# Improved Python Package for DNA Sequence Encoding using Frequency Chaos Game Representation

**DOI:** 10.1101/2024.04.14.589394

**Authors:** Abhishek Halder, Piyush, Bernadette Mathew, Debarka Sengupta

## Abstract

**Summary:** Frequency Chaos Game Representation (FCGR), an extended version of Chaos Game Representation (CGR), emerges as a robust strategy for DNA sequence encoding. The core principle of the CGR algorithm involves mapping a onedimensional sequence representation into a higher-dimensional space, typically in the two-dimensional spatial domain. This paper introduces a use case wherein FCGR serves as a kmer frequency-based encoding method for motif classification using a publicly available dataset.

**Availability and implementation:** The FCGR python package, use case, along with additional functionalities, is available in the [GitHub](https://github.com/abhi1238/FCGR). Our FCGR package demonstrates superior accuracy and computational efficiency compared to a leading R-based FCGR library [1], which is designed for versatile tasks, including proteins, letters, and amino acids with user-defined resolution. Nevertheless, it is important to note that our Python package is specifically designed for DNA sequence encoding, where the resolution is predetermined based on the kmer length. It is a drawback of our current package compared to the state-of-the-art R-based kaos package [1].

## Introduction

The CGR algorithm, initially designed for making fractals [2], was later adapted by Jeffrey to analyze DNA sequences [3]. The technique, rooted in chaotic dynamics, creates a visual representation of gene sequences with detailed structures, capturing both local and global patterns [3].

Here, we introduce an enhanced version of the R kaos package [1], featuring significantly improved processing speed and precise outcomes. Additionally, the current kaos Python package empowers a broad community of Python researchers by simplifying the generation of the FCGR matrix.

### Improvements and new features

#### Accurate calculation of FCGR matrix

The existing R kaos package [1], which relies on a set of predefined equations to generate FCGR matrices by constructing n-flakes polygons, may not consistently provide accurate results. We have replaced this approach with a more efficient and optimized algorithm that accurately counts the frequency of each kmer, ensuring precise numbers for each kmer.

#### Time efficiency

The current state-of-the-art R kaos package [1] is considerably slower when compared to the Python-based kaos package. Our efficient and optimized algorithm offers significantly faster execution speeds than the R-based kaos package.

#### Introduction of new functionality

In addition to the standard generation of FCGR encodings, this paper introduces enhanced functionalities for users. These include the ability to determine the frequency of a specific kmer, retrieve the index of a particular kmer in the FCGR encoding, search for a kmer at a specified index, and obtain a dictionary containing the frequency of each kmer. These added functionalities significantly enhance the analytical capabilities for users utilizing the current Python package. The details of all these functionalities are outlined in the [documentation](https://github.com/abhi1238/FCGR/blob/main/package/kaos%20documentation.pdf) of the GitHub repository.

### FCGR formulation

#### Chaos Game Representation for triangle

The CGR was initially used for the generation of the Sierpinski triangle Fig. 1. As shown in Fig. 2, the process begins by selecting a random starting point (S) within the Sierpinski triangle. A vertex (A) is then randomly chosen, and a point (P1) is plotted halfway to that vertex (A). This procedure continues, with P1 becoming the new starting point. The second point (P2) is drawn halfway to another randomly chosen vertex (B). The trajectory of the walker is shown in the Fig. 2. Through the iterative application of this algorithm, the Sierpinski triangle is progressively formed, as illustrated in Fig. 1 [4].

**Figure 1:**
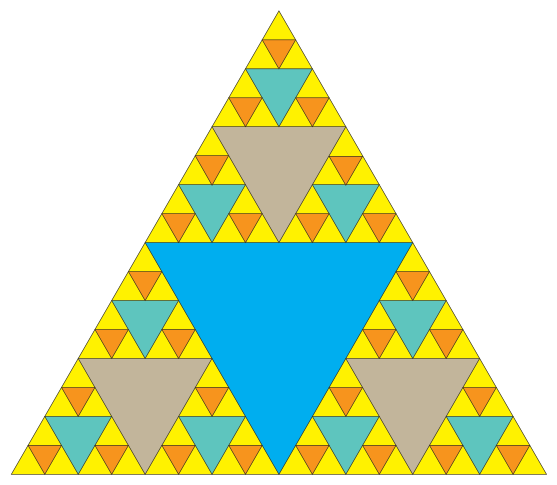
Sierpinski triangle

**Figure 2:**
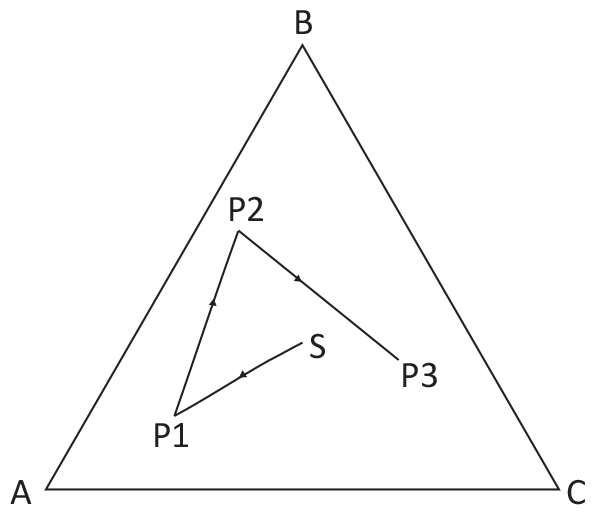
Walker trajectory inside Sierpinski triangle

#### Chaos Game Representation for squares and its role in DNA encoding

Instead of using a triangle, the CGR for DNA was based on a square, with the four vertices representing the four nucleotides, adenine (A), cytosine (C), guanine (G), and thymine (T), as shown in the Fig. 3. Fig. 3 depicts the trajectory of the walker utilizing the chaos game representation with four vertices. The rules for the movement of the walker inside the square, starting from S, are the same as those mentioned for the triangle case in the above section.

**Figure 3:**
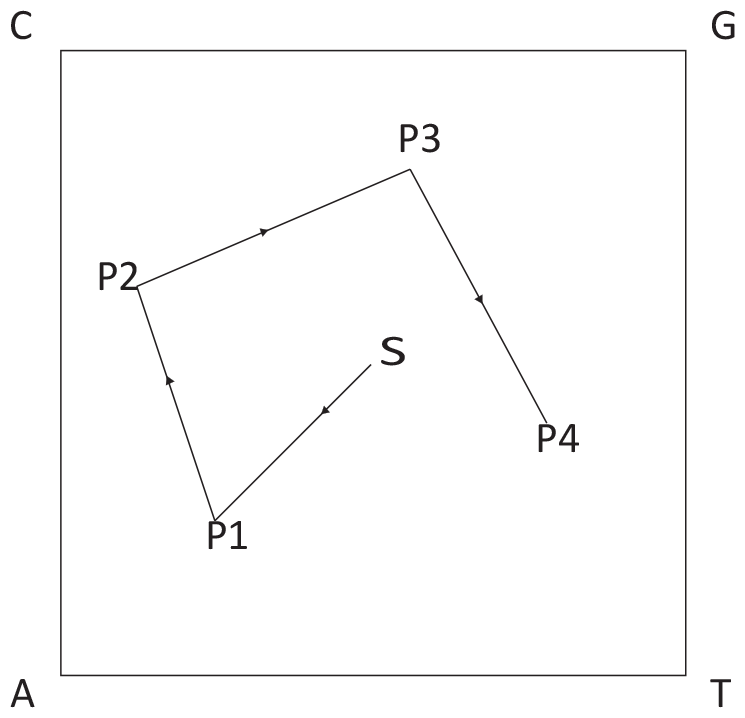
CGR algorithm walker trajectory with four vertices

#### Mathematical formulation of CGR for DNA

Let *S* be a DNA nucleotide string of length *N*, where *S*[*i*] represents the *i*-th symbol (1 ≤*i* ≤*N*) corresponding to a DNA nucleotide (A, C, G, T).

The notation *S*[..*i*] denotes the DNA sequence prefix ending at position *i* (*S*[..*i*] = *S*[1..*i*]). The CGR iterative algorithm operates on ℝ ^2^, with each vertex corresponding to a DNA base (A, C, G, T). For a given DNA sequence *S* of length *N*, CGR maps each *S*[..*i*] prefix to the point *x*_*i*_ ∈ ℝ ^2^ using an iterative procedure. It can be written in equation form as given below,

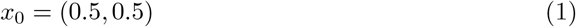

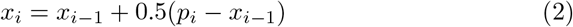

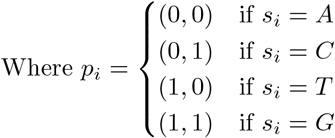

In the original formulation [3], the initial point *x*_0_ was taken as the center of the square (0.5, 0.5). Alternatively, this point could be chosen randomly within the square [5]. The user can also customize the vertex for *p*_*i*_ corresponding to a given *s*_*i*_. Here, *s*_*i*_ represents the randomly chosen vertex for the *i*^*th*^ step of the walker.

### Frequency Chaos Game Representation Matrix

A significant application of CGR involves assessing kmer abundance through FCGR [3, 6]. FCGR, a simplified version of CGR, reduces noise by counting points on a grid applied to the CGR space, creating a matrix that visualizes the frequency of kmers in a predefined order, as shown in Fig. 4. This approach aids in identifying patterns and similarities in genetic information. The number of quadrants in an FCGR grid can be calculated as 4^*k*^, where k denotes the kmer size. The FCGR matrix dimension is 2^*k*^ X 2^*k*^.

**Figure 4:**
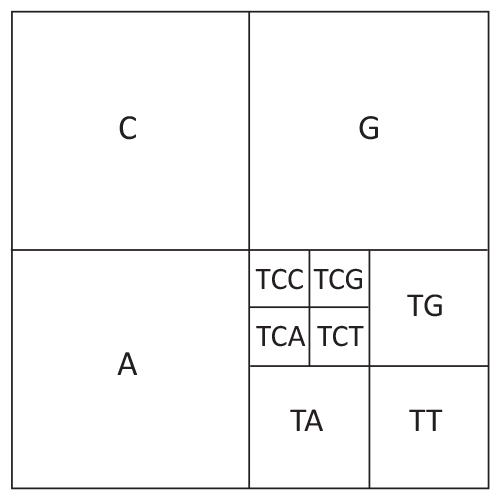
Division of the CGR space due to the iterative process [4]

Jeffrey [3] observed that while random sequences lack meaningful patterns, the application of CGR to DNA sequences reveals fractal patterns, signifying non-random structures. The Fig. 5 illustrates the heatmap of *H. sapien* DNA in FCGR-encoded format (negative log scale after frequency normalization).

**Figure 5:**
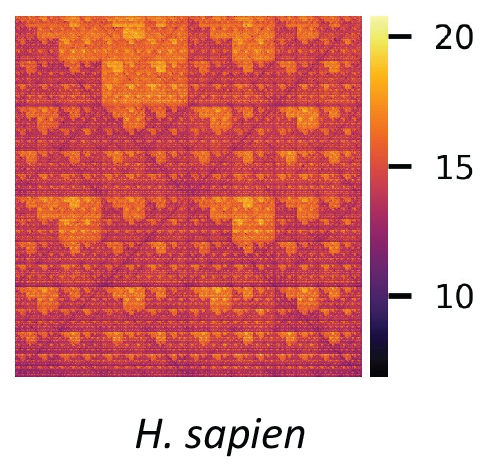
Heatmap of *H. sapien* DNA FCGR

The presence of repetitive fractal patterns in the FCGR of DNA is evident, as depicted in Fig. 5, supporting the findings of Jeffrey [3].

### Key steps involved in FCGR encoding

The initial step involves the user reading the FASTA file using the **read fasta** function. It is important to note that the read fasta method concatenates all the contigs in the same order as they appear in the FASTA file. It is advisable to use the whole genome sequence when creating the FCGR matrix from a FASTA file to ensure the integrity of the DNA sequence. Following this, the FASTA obtained in the preceding step serves as the input for the **Chaos frequency matrix** function. This function provides the frequency of each kmer in the form of a chaos game representation. All these functions are accessible within the Python kaos package.

### Comparative results

In Fig 6, the correlation analysis shows how the FCGR values of *E. coli* ^1^ DNA from the R kaos package [1] varies compared to those from the Python kaos package introduced in this paper. As the length of the kmer increases, the correlation between the two packages’ FCGR values decreases. We emphasize that our proposed package calculates FCGR using the actual frequency count of kmers, ensuring the accuracy of the resulting FCGR matrix. It is evident from Figure 6 that the accuracy of the FCGR matrix obtained from the R-based kaos package [1] decreases significantly as the kmer length increases.

**Figure 6:**
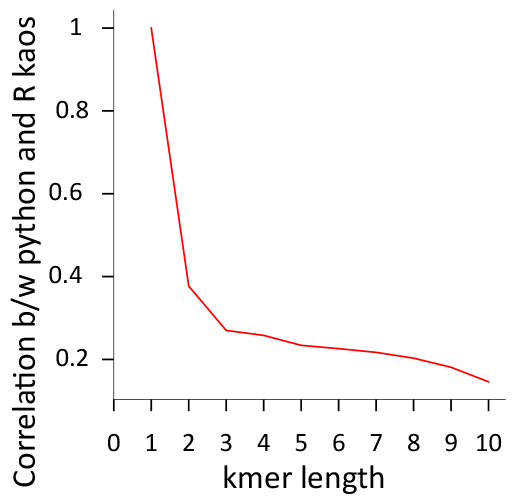
FCGR correlation plot between R kaos [1] and current package for *E. coli* DNA

Additionally, we conducted a comparison of the computational efficiency of both packages. Notably, our Python package showed much faster processing times than the R kaos package [1], as illustrated in Fig 7 ^2^.

**Figure 7:**
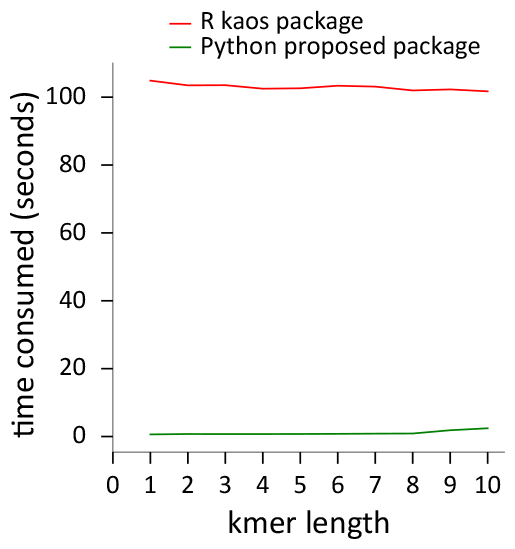
Time taken by R kaos package [1] and current package for *E. coli* DNA

However, it’s important to note that our newly proposed Python kaos package is specifically tailored for DNA sequences. In contrast, the R kaos package by Lochel et al. [1] offers versatility by accommodating various data types, such as protein sequences and letters. Furthermore, the R kaos package supports user-defined resolution, a feature not currently implemented in our Python version, which allows users to set resolutions according to their specific needs. This versatility offers advantages for a variety of applications.

### Usecase of FCGR

#### Dataset

Within DNA there are repeats known as motifs. Sequence motifs are short, recurring patterns in DNA that are presumed to have a biological function. Detecting these motifs, which are shared among a group of functionally related sequences, is crucial for understanding the genome and its operations. The dataset utilized to predict protein binding sites was sourced from a previous paper [7]. This dataset comprises artificially generated DNA sequences, each of length 50 bases. Every nucleotide sequence is paired with a label, signifying whether the specific DNA sequence is associated with a protein binding site in DNA. Within the provided set of 2000 nucleotide sequences, 987 sequences were identified as protein binding sites, while the remaining sequences denote instances of non-protein binding sites.

#### Methodology

The details of the data preprocessing, the architecture of the deep learning model, and the data splitting strategy are outlined in Algorithm 1.

##### Algorithm 1

Algorithm for protein binding site prediction

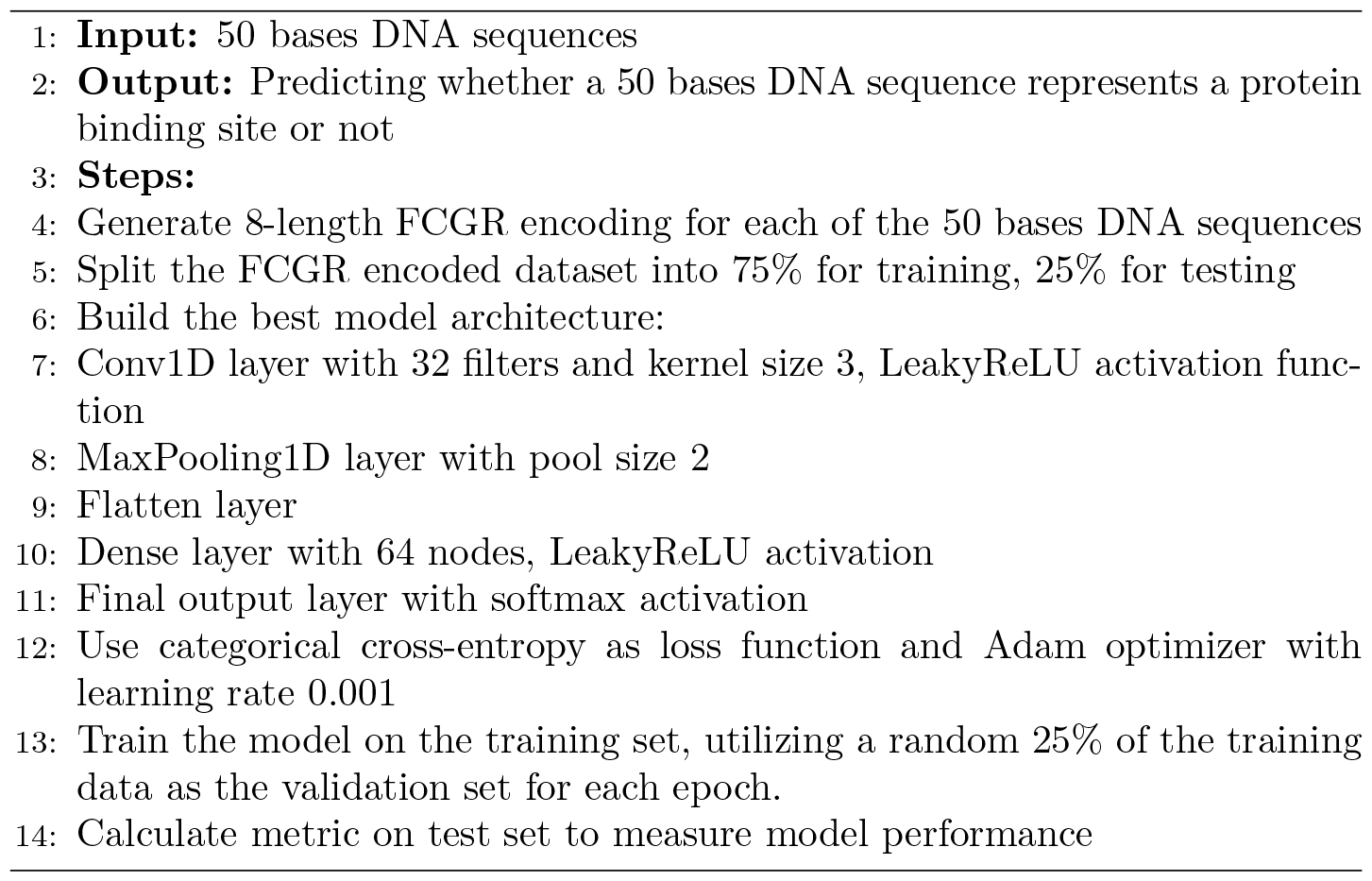

#### Evaluation metrics

The model is evaluated based on a few metrics such as accuracy, precision, recall, F1 score, and Cohen’s Kappa scores. Accuracy represents the proportion of correctly classified instances among the total number of instances. Precision is the ratio of correctly predicted positive observations to the total predicted positives. Recall is the ratio of correctly predicted positive observations to all actual positives. The F1 score is the harmonic mean of precision and recall, providing a balance between the two metrics. Cohen’s Kappa is a statistic that measures inter-rater agreement for categorical items.

## Results

The evaluation metrics on the test set, including accuracy, precision, recall, F1 score, and Cohen’s kappa, are recorded as 0.998, 0.996, 1, 0.997, and 0.996, respectively.

### Comparision of FCGR encoding with the one hot encoding

As demonstrated in the [Google Colab file](https://colab.research.google.com/drive/17E4h5aAOioh5DiTo7MZg4hpL6Z_0FyWr#scrollTo=J1bvxV9J-EMT) provided by the author in the previous study [7] on motif finding, the use of the one-hot encoding technique for DNA sequence representation resulted in a test set accuracy of 98%. In contrast, our proposed method, which utilizes FCGR for encoding, achieved an accuracy of 99.8% on the same test dataset. While the performance improvement may not be large, it highlights the effectiveness of encoding strategies using DNA sequence information, hinting at potential benefits for model learning.

## Footnotes

1 the FASTA of *Escherichia coli* str. K-12 substr. MG1655 is obtained from the [NCBI](https://www.ncbi.nlm.nih.gov/datasets/genome/GCF_000005845.2/)

2 The FCGR matrix generation time is reported based on an 11th Gen Intel(R) Core(TM) i7-11700 @ 2.50GHz system, with 96 GB RAM and a 1TB Solid State Drive, operating on Ubuntu 22.04.2 LTS

